# Supramodal representation of the sense of body ownership in the human parieto-premotor and extrastriate cortices

**DOI:** 10.1101/2022.08.28.502233

**Authors:** Yusuke Sonobe, Toyoki Yamagata, Huixiang Yang, Yusuke Haruki, Kenji Ogawa

## Abstract

The sense of body ownership, defined as the sensation that one’s body belongs to oneself, is a fundamental component of bodily self-consciousness. Several studies have shown the importance of multisensory integration for the emergence of the sense of body ownership, together with the involvement of the parieto-premotor and extrastriate cortices in bodily awareness. However, whether the sense of body ownership elicited by different sources of signal, especially visuotactile and visuomotor inputs, is represented by common neural patterns remains to be elucidated. We used functional magnetic resonance imaging (fMRI) to investigate the existence of neural correlates of the sense of body ownership independent of the sensory modalities. Participants received tactile stimulation or executed finger movements while given synchronous and asynchronous visual feedback of their hand. We used multi-voxel patterns analysis (MVPA) to decode the synchronous and asynchronous conditions with cross-classification between two modalities: the classifier was first trained in the visuotactile sessions and then tested in the visuomotor sessions and vice versa. Regions of interest-based and searchlight analyses revealed significant above-chance cross-classification accuracies in the bilateral intraparietal sulcus (IPS), the bilateral ventral premotor cortex (PMv), and the left extrastriate body area (EBA). Moreover, we observed a significant positive correlation between the cross-classification accuracy in the left PMv and the difference in subjective ratings of the sense of body ownership between the synchronous and asynchronous conditions. Our findings revealed the neural representations of the sense of body ownership in the IPS, PMv, and EBA that is invariant to the sensory modalities.

**Significance Statement:** Previous studies have shown neural correlates of the sense of body ownership in parieto-premotor and extrastriate cortices. However, whether the sense of body ownership induced by different sensory inputs is represented in common neural patterns remains unelucidated. Using functional magnetic resonance imaging (fMRI) with multi-voxel pattern analysis (MVPA), we investigated neural representations of the sense of body ownership invariant to modalities. Decoding neural patterns for visuotactile and visuomotor modalities revealed successful cross-classification accuracies in intraparietal sulcus (IPS), ventral premotor cortex (PMv), and extrastriate body area (EBA). Furthermore, cross-classification accuracy in PMv was positively correlated with subjective ratings of the sense of body ownership. These findings demonstrate that supramodal representations in parieto-premotor and extrastriate cortices underlie the sense of body ownership.

## Introduction

The brain receives multiple signals from the sensory channels. The integration of multimodal sensory information contributes to the coherent representation of the self (Tsakiris, 2010, 2017; Petkova et al., 2011; Blanke, 2012). Previous research in macaques has empirically revealed the involvement of the parieto-premotor cortices in the process of multisensorial information about the body vicinity. Neurons in the premotor and the parietal cortices are modulated by more than one sensory signal, and the integration is maximized when stimuli are spatiotemporally congruent. Considering that multisensory integration requires different sensory inputs to be unified in individual neurons, bimodal neurons are ideal candidates for multisensory convergence (Graziano et al., 1997; Duhamel et al., 1998; Avillac et al., 2007).

The sense of body ownership is a component of bodily self-consciousness based on multisensory integration and is defined as the sensation that part of or the whole body belongs to oneself. It has been approached by theoretical papers from both a phenomenological (Gallagher, 2000; Blanke and Metzinger, 2009) and a neurocognitive (Tsakiris, 2010; Blanke, 2012) perspective. Theoretical and empirical work has investigated the sense of body ownership, using the rubber hand illusion (RHI) (Armel and Ramachandran, 2003; Tsakiris and Haggard, 2005; Apps and Tsakiris, 2014; Tsakiris, 2017). RHI is an experimental paradigm, whereby participants feel ownership of a fake hand placed in front of them after synchronous touch of their real hand and the fake one (Botvinick and Cohen, 1998). Converging evidence suggests that the illusion is induced when visual and proprioceptive signals are synchronized (Shimada et al., 2005, 2009, 2014; Bekrater–Bodmann et al., 2014).

RHI has been widely employed to investigate multisensory integration and the sense of body ownership albeit with methodological differences (for review, see Riemer et al., 2019). This might raise the issue of whether multisensory integration under different conditions induces the same perceptual experience. Although the findings are not consistent and depend on whether subjective ratings or proprioceptive drift were utilized, previous behavioral studies involving a direct comparison of visuotactile and visuomotor conditions reported that both conditions result in a similar ownership sensation (Tsakiris et al., 2006; Kalckert and Ehrsson, 2014a). However, whether the behaviorally similar sense of body ownership is represented by a common or by distinct spatial neural activities is unknown. Neuroimaging studies have identified brain regions associated with the sense of body ownership, such as the intraparietal sulcus (IPS), the ventral premotor cortex (PMv), and the extrastriate body area (EBA). However, the neural correlates of the sense of body ownership induced by tactile stimulation and kinesthetic movement have been investigated independently by different studies (Ehrsson et al., 2004; Tsakiris et al., 2010; Gentile et al., 2011; Olivé et al., 2015; Lee and Chae, 2016; Limanowski and Blankenburg, 2016b). Furthermore, prior functional magnetic resonance imaging (fMRI) studies have mainly conducted a univariate analysis of the overall activation increase in a region, not providing information about the similarities in the spatial activation patterns between different sensory modalities.

Here, we aimed to examine whether the sense of body ownership induced by visuotactile and visuomotor inputs is represented in common neural activation patterns. We manipulated the participant’s sense of ownership of their hand using tactile stimulation and finger movement within a single fMRI experiment. We then used multi-voxel pattern analysis (MVPA). In contrast to the conventional subtraction-based fMRI analysis, MVPA enables the identification of differences in spatial patterns of activated voxels in a region (Haynes and Rees, 2005; Kamitani and Tong, 2005; Kriegeskorte et al., 2006; Norman et al., 2006). We thus examined whether differences in the multi-voxel spatial patterns in synchronous and asynchronous conditions were common to each modality and predicted that the IPS, PMv, and EBA, regions associated with the sense of body ownership, could decode the differences regardless of the modalities.

## Materials and Methods

### Participants

Twenty-six healthy volunteers (14 males and 12 females) with a mean age of 22.38 years (20–28 years) participated in the experiment. The number of participants was determined based on previous fMRI experiments (Gentile et al., 2011; Bekrater–Bodmann et al., 2014). All participants were right-handed as assessed by a modified version of the Edinburgh Handedness Inventory (Oldfield, 1971). Written informed consents were obtained from all participants in accordance with the Declaration of Helsinki. The experimental protocol was approved by the local ethics committee. All participants were naïve to the purpose of the fMRI experiment, although two of them had previously participated in behavioral RHI experiments. One participant with large head movements (maximum translation per session, 66.44 mm) during the scanning was excluded from the analysis. Thus, the final analysis included 25 participants.

### Experimental design and statistical analysis

We employed a two-by-two within-subject factorial design. We manipulated the type of modalities (i.e., visuotactile vs. visuomotor) and the timings of the visual feedback (i.e., synchronous vs. asynchronous). In the visuotactile condition, the participant’s right index finger was stroked with a paintbrush. In the visuomotor condition, the participants raised and lowered their right index finger on their own. In the synchronous condition, videos of the strokes or movements were presented in real-time. In the asynchronous condition, the videos were presented with a 1000 ms delay. The four conditions were labeled as visuotactile synchronous (TS), visuotactile asynchronous (TA), visuomotor synchronous (MS), and visuomotor asynchronous (MA).

Statistical analyses were performed using a two-way within-subject analysis of variance (ANOVA), a one-sample *t*-test, and a paired *t*-test. To control for the problem of multiple comparisons, we applied the Holm–Bonferroni procedure (Holm, 1979) based on the number of regions of interest (ROIs) in the left and right hemispheres, respectively. We used Pearson’s *r* to investigate the correlation between the cross-classification accuracies and the subjective ratings in the questionnaire. The statistical significance level is reported for each analysis.

### Experimental setup

The participants lay comfortably in a supine position on the bed in the MRI scanner. The participant’s hand was extended on the bed in a relaxed position and hidden from their own view. A flexible arm (Articulated arm mount, MRC Systems GmbH, Heidelberg, Germany) was attached to the head coil with an MRI-compatible color camera (12M-i, MRC Systems GmbH, Heidelberg, Germany) fixed at the end of the arm. This camera captured videos of the participant’s right hand. The videos were projected on the monitor using Light Capture (I-O DATA, Ishikawa, Japan) in the control room in an anatomically congruent frame with respect to the real hand. A webcam (C270n, Logitech, Lausanne, Switzerland) captured the videos on the monitor using MATLAB (The MathWorks, Inc., MA, USA). The videos were presented on a liquid crystal display (NNL LCD Monitor, NordicNeuroLab, Bergen Norway) and projected onto a custom-made viewing screen. The participants viewed the screen via a mirror. Real-time video transfer was realized with a minimal intrinsic delay of approximately 400 ms in the synchronous condition. The intrinsic delay refers to the time difference between the movement of the right hand and the movement of the hand on the screen. It was calculated by assessing how long (in ms) after the movement of the real hand, the movement of the hand appeared on the LCD using Adobe Premiere Pro (Adobe Inc., CA, USA). In the asynchronous condition, a systematic delay of 1000 ms was added to the intrinsic delay. The distance between the camera and the real hand was adjusted before the experiment according to the size of the participant’s hand to make it easier for the participant to see their hand on the screen via a mirror. The video presentation was implemented by Computer Vision Toolbox (https://mathworks.com/products/computer-vision.html), Image Processing Toolbox (https://mathworks.com/products/image.html), MATLAB Support Package for USB Webcams (https://mathworks.com/matlabcentral/fileexchange/45182-matlab-support-package-for-usb-webcams), and Psychtoolbox (http://psychtoolbox.org/) for MATLAB.

Participants heard a beep through MRI-compatible headphones (Resonance Technology Inc., CA, USA) to ensure that the number of finger movements was the same among participants. They heard the beep both in the visuotactile and visuomotor sessions to prevent a confounding effect of the beep. The beep sound was created at a frequency of approximately 1 Hz with a jitter of −200 ms, 0 ms, or +200 ms added pseudo-randomly to make the stimulation sequence unpredictable. The beep output from the computer was split in two: one output to the participant’s headphones and another sounded in the entire MRI room via the intercom as a cue for the experimenter to apply brushstrokes to the participant’s hand. The beep sounded in the entire MRI room both in the visuotactile and visuomotor sessions. The beep presentation was implemented by MATLAB (Figure 1).

**Figure 1:**
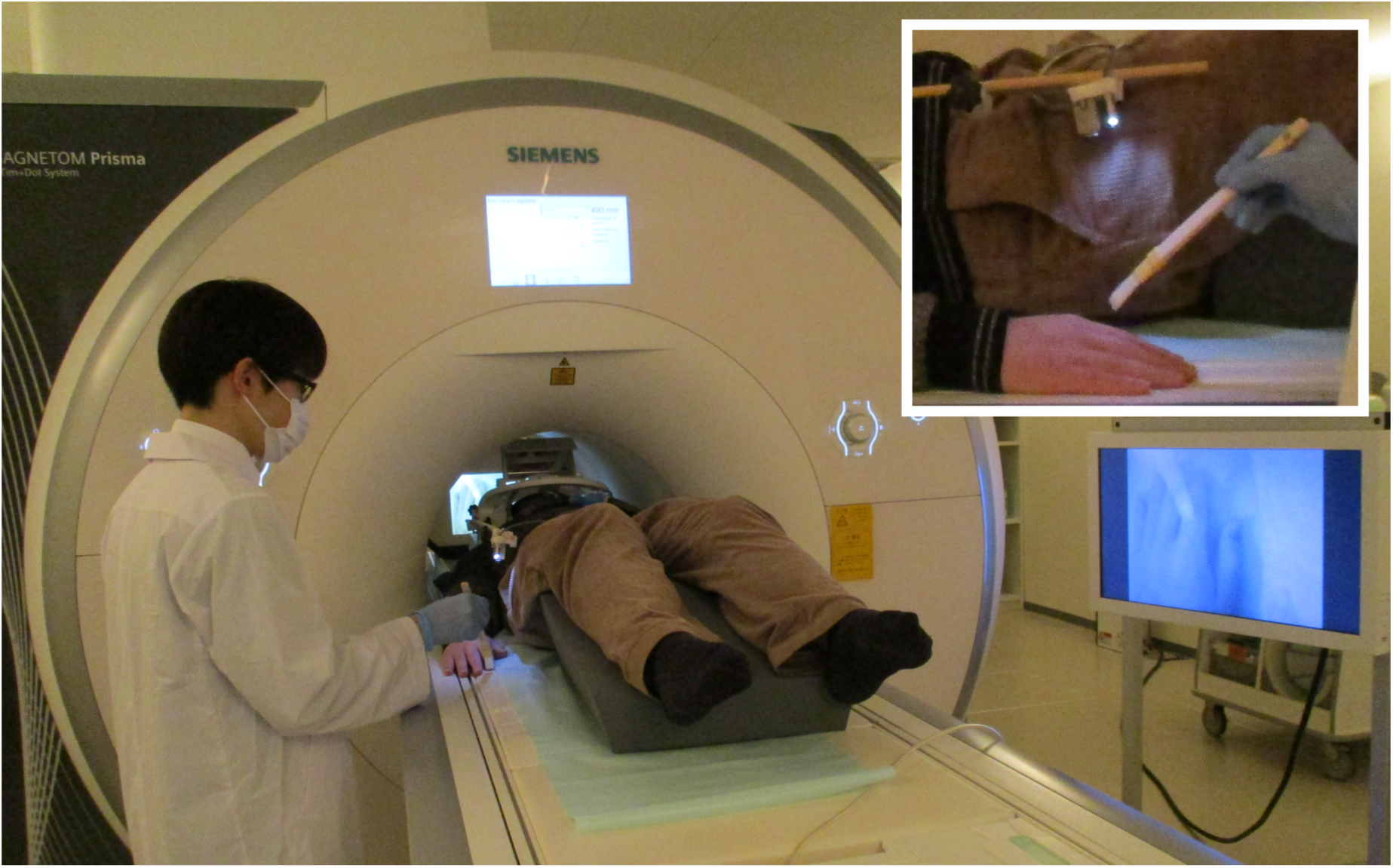
Experimental setup. The participants watched videos of their right hand captured by the camera attached to the end of the flexible arm on the head coil. In the visuotactile sessions, an experimenter stroked the participant’s right index finger at a frequency of approximately 1 Hz to a beep sounded in the MRI room. In the visuomotor sessions, the participants lifted and lowered their right index finger on their own at the same rate to a beep heard through headphones while an experimenter stood beside them.

### Task procedures

There were four experimental sessions. The visuotactile and visuomotor sessions were each performed twice in succession. The order of sessions was pseudo-randomized across the participants. Each session consisted of 16 trials, 8 in synchronous and 8 in asynchronous conditions. Synchronous or asynchronous conditions were determined pseudo-randomly, but the same type of visual feedback was not repeated three times in succession.

Each trial began with 2 s of yellow fixation cross, followed by 18 s of videos of the participant’s right hand. In the visuotactile sessions, tactile stimulation was delivered by an experimenter manually with a paintbrush at a frequency of approximately 1 Hz during 18 s, as cued by a beep. The upper part of the participant’s right index finger was stroked from the fingertip to the knuckle at one beep and inversely from the knuckle to the fingertip at the next beep. To know when to start and end stroking, an experimenter checked the LCD placed adjacent to the MRI scanner during the stimulation. Because of a malfunction of the projector, the LCD was placed near the participant’s head for five subjects. The five participants viewed the videos on the LCD via a mirror. In these cases, an experimenter stroked the participant’s right index finger while looking inside the MRI scanner to check the LCD. In the visuomotor sessions, the participants lifted their right index finger at one beep and lowered it at the next beep at the same rate during 18 s. To remove the confounding influence of the presence of people between the visuotactile and visuomotor sessions, an experimenter kept standing beside the participant in the same position as in the visuotactile sessions. Subsequently, a white fixation cross was presented for 8 s and served as baseline (Figure 2). The participants heard a beep via MRI-compatible headphones throughout the fMRI scanning period. The videos were recorded and analyzed after the experiment. The average (SD) number of brush strokes and finger movements for each participant was 16.55 (0.95). The participants practiced the finger movements for one minute before they entered the MRI scanner.

**Figure 2:**
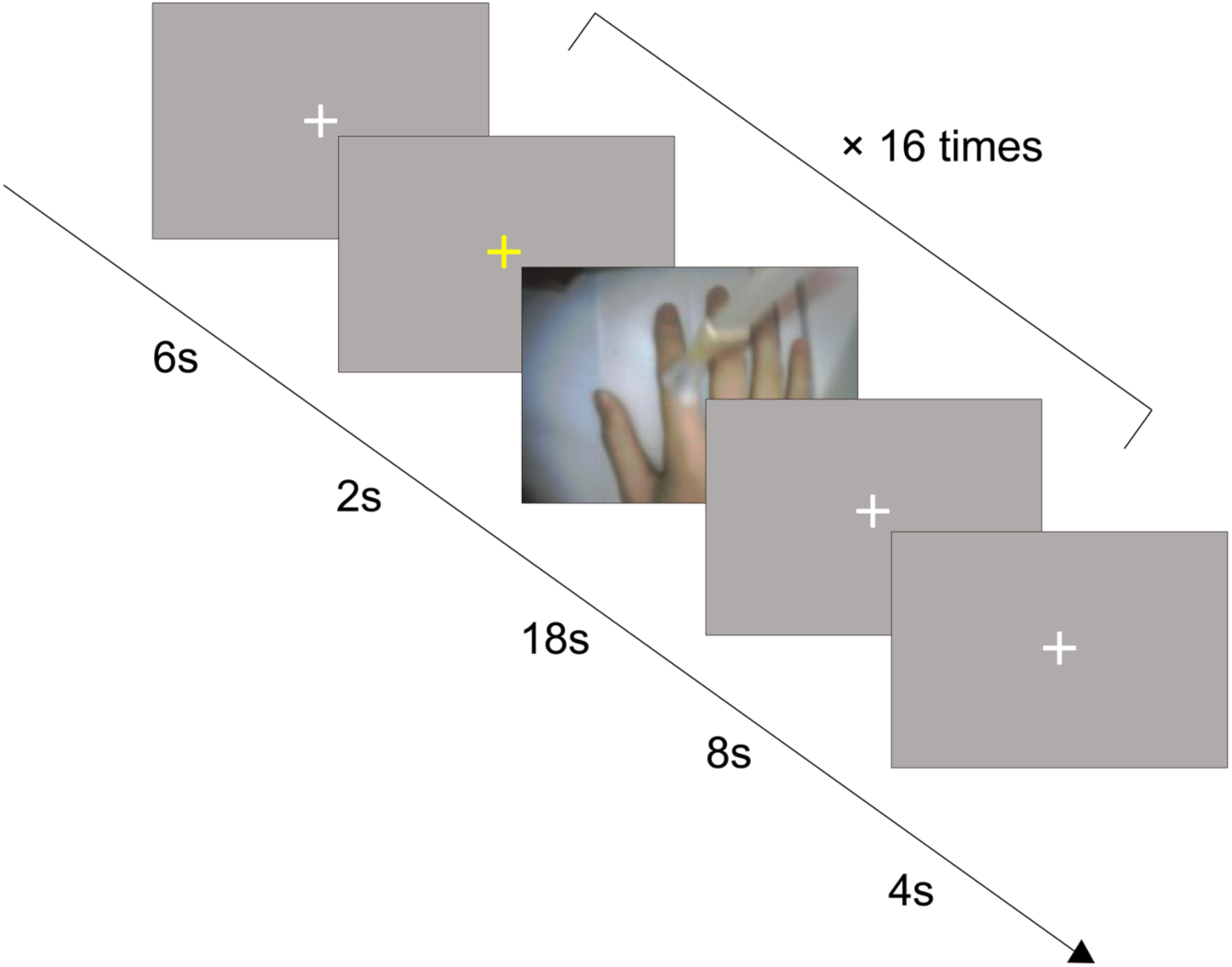
Schematic depiction of the time course of the visuotactile and visuomotor sessions. Trials began with the yellow fixation cross as the cue. Two seconds later, the videos of the right hand were displayed for 18 s, followed by the white fixation cross for 8 s. In 8 out of 16 trials, the videos were presented synchronously with the tactile stimuli or finger movements. In the other 8 trials, the videos were displayed with a systematic delay of 1000 ms. The videos were never synchronous or asynchronous three times in succession.

After every two sessions, the participants completed questionnaires to report their subjective experience in the MRI scanner (Table 1). The questionnaire items were based on those used by Kalckert and Ehrsson (2014a) and Kalckert and Ehrsson (2014b). The questionnaire consisted of six items with items (1)–(4) serving as indicators of the sense of body ownership (Ownership) and with items (5) and (6) serving as controls for ownership, in particular for task compliance, suggestibility, and expectancy effects (Ownership control) (Kalckert and Ehrsson, 2014a, 2014b). The item order was pseudo-randomized. Responses were self-paced and were made using a 7-point Likert scale, ranging from +3 (strongly agree) to −3 (strongly disagree), with 0 indicating neither agreement nor disagreement or “uncertainty.” We recorded the responses using MRI-compatible response pads (Current Design, Philadelphia, USA). The participants answered the six items twice in succession, once for the synchronous condition and once for the asynchronous condition. The order of questionnaire items was pseudo-randomized across the participants, but with the same order across the modalities within a participant. To avoid the confounding effect of the hands used to rate items, the participants were pseudo-randomly divided into two groups according to which hand they used to rate items as positive. The right index finger, which was stimulated by the paintbrush and used for finger movements, was not assigned to press the button. The questionnaire presentation was implemented by PsychoPy (Peirce et al., 2019).

**Table 1:** Statements used in the questionnaire to quantify the subjective experience of the sense of body ownership

|  |  |  |
| --- | --- | --- |
| Ownership | 1 | I felt as if I was looking at my own hand |
|  | 2 | I felt as if the hand in the video was part of my body |
|  | 3 | I felt as if the hand in the video was my hand |
|  | 4 | I felt as if my real hand was turning the hand into the video |
| Ownership control | 5 | It seems as if I had more than one right hand |
|  | 6 | It felt as if I had no longer a right hand, as if my right hand had disappeared |

As the present study did not use a fake hand but presented their own hand to the participants via video feedback, the item “I felt as if my real hand was turning the hand into the video” adapted from Kalckert and Ehrsson (2014a) and Kalckert and Ehrsson (2014b) was analyzed as Ownership.

### Functional EBA localizer scan

An independent functional localizer scan was performed for all participants after the visuotactile and visuomotor sessions to determine a ROI of the EBA selectively activated by body parts (Downing et al., 2001). The protocol was adapted from a previous study (Taylor et al., 2007). A total of 16 blocks were performed, which consisted of 8 body blocks and 8 control blocks. A body block consisted of 20 images of headless human body parts in different postures, which were alternated with a control block consisting of 20 images of chairs. All images were grayscale. Each image was presented for 300 ms, followed by a black screen for 450 ms. A fixation cross was intercalated at the end of each block for 12 s as baseline. To maintain and monitor attention, the participants performed a one-back repetition detection task during the scan. The same images were presented twice in succession, twice during each block. The participants were asked to press a button with the right index finger when they detected the immediate repetitions. The responses were recorded using MRI-compatible response pads. The participants practiced a shorter version of the task outside the MRI scanner before the experiment. The stimuli presentation was implemented by PsychoPy (Peirce et al., 2019).

### MRI data acquisition

All scans were performed on a Siemens (Erlangen, Germany) 3-Tesla Prisma scanner with a 20-channel head coil at Hokkaido University. T2*-weighted echo planar imaging (EPI) was used to acquire a total of 229 and 219 scans per session for the main and localizer sessions, respectively, with a gradient echo EPI sequence. The first three scans of each session were discarded to allow for T1 equilibration. The scanning parameters were repetition time (TR), 2,000 ms; echo time (TE), 30 ms; flip angle (FA), 90°; field of view (FOV), 192 × 192 mm; matrix, 94 × 94; 32 axial slices; and slice thickness, 3.5 mm with a 0.875 mm gap. T1-weighted anatomical imaging with an MP-RAGE sequence was performed using the following parameters TR, 2,300 ms; TE, 2.32 ms; FA, 8°; FOV, 240 × 240 mm; matrix, 256 × 256; 192 axial slices; and slice thickness, 0.90 mm without a gap.

### Processing of fMRI data

Image preprocessing was performed using the SPM12 software (Wellcome Department of Cognitive Neurology, http://www.fil.ion.ucl.ac.uk/spm). All functional images were initially realigned to adjust for motion-related artifacts. Volume-based realignment was performed by co-registering images using rigid-body transformation to minimize the squared differences between volumes. The realigned images were then spatially normalized with the Montreal Neurological Institute (MNI) template based on the affine and non-linear registration of co-registered T1-weighted anatomical images (normalization procedure of SPM). They were resampled into 3-mm-cube voxels with sinc interpolation. Images were spatially smoothed using a Gaussian kernel of 6 × 6 × 6 mm full width at half-maximum. However, images used for MVPA were not smoothed to avoid blurring the fine-grained information contained in the multi-voxel activity (Mur et al., 2009; Kamitani and Sawahata, 2010).

Using the general linear model, the 16 blocks per session were modeled as 16 separate boxcar regressors that were convolved with a canonical hemodynamic response function. Low-frequency noise was removed using a high-pass filter with a cut-off period of 128 s, and serial correlations among scans were estimated with an autoregressive model implemented in SPM12. This analysis yielded 16 independently estimated parameters (beta values) per session for each individual voxel. These parameter estimates were then *z*-normalized across voxels for each trial and were subsequently used as inputs to MVPA.

### Definition of ROIs

We chose the IPS and PMv as ROIs because previous studies revealed that these regions are primarily involved in the multisensory processes implicated in the sense of body ownership. Electrophysiological studies reported that neurons located in the premotor and parietal cortices of macaques respond to stimuli from not only one but also multiple sensory modalities (Graziano et al., 1997; Duhamel et al., 1998; Avillac et al., 2007). This finding is supported by neuroimaging research in humans revealing greater activations in the IPS and PMv when both visual and tactile stimuli are presented compared with the activation resulting from either a visual or a tactile stimulus (Gentile et al., 2011). Studies using RHI consistently found that multisensory integration occurs in the IPS and PMv when stimuli are spatiotemporally congruent, resulting in the self-attribution of a fake hand (Ehrsson et al., 2004; Limanowski and Blankenburg, 2015; Olivé et al., 2015). Recent studies using transcranial magnetic stimulation (TMS) and transcranial direct current stimulation (tDCS) also revealed the involvement of these regions in the sense of body ownership (Karabanov et al., 2017; Convento et al., 2018; Lira et al., 2018). Therefore, the IPS and PMv are likely responsible for the multisensorial process of body ownership. The bilateral IPS was defined using Anatomy toolbox (Eickhoff et al., 2005), and the bilateral PMv was defined using the Human Motor Area Template (Laboratory for Rehabilitation Neuroscience, http://lrnlab.org/; Mayka et al., 2006).

Additionally to the frontoparietal regions, the EBA, which is part of the lateral occipital cortex (LOC), was chosen as ROI. The EBA has been associated with the visual processing of parts of or all the human body by Downing et al. (2001). However, recent neuroimaging studies have shown the involvement of the EBA in the sense of body ownership. For example, Limanowski and Blankenburg (2015) revealed increased connectivity between the LOC and IPS during RHI. The EBA activity is positively correlated with subjective illusion scores (Limanowski et al., 2014). Wold et al. (2014) reported an increased proprioceptive drift toward the rubber hand after rTMS of the left EBA. The EBA cannot be identified using templates based on anatomical structures. Using a localizer task is one approach to solve the problem. We measured a significant activation of the EBA during the presentation of body part images compared with that elicited by chair images and determined a sphere with a radius of 5 mm centered on MNI coordinates of the group-level analysis with a threshold of *p* < 0.05 corrected for family-wise error (FWE) with an extent threshold of 10 voxels (MNI coordinates: [−48, −70, 8] for the left hemisphere and [51, −58, 2] for the right hemisphere). A similar approach was used by Olivé et al. (2015).

### Mass-univariate analysis

We used the conventional mass-univariate analysis of individual voxels to reveal areas activated in each condition and its combination. Firstly, we analyzed the main effect of modalities collapsed across timings of visual feedback, [(TS + TA) − (MS + MA)] and inversely [(MS + MA) − (TS + TA)]. Secondly, we analyzed the main effect of timings of visual feedback collapsed across modalities, [(TS + MS) − (TA + MA)] and inversely [(TA + MA) − (TS + MS)]. Thirdly, we analyzed the interaction effects between modalities and timings of visual feedback, [(TS−TA) − (MS−MA)] and inversely [(MS−MA) − (TS−TA)]. For the analysis of the EBA localizer scan, we compared areas activated during the presentation of body parts with regions activated by chair pictures.

Contrast images were generated for each participant using a fixed-effects model and were analyzed using a random-effects model of a one-sample *t*-test. Activation was reported with a threshold of *p* < 0.05 corrected for FWE with an extent threshold of 10 voxels. If no area survived the threshold, activations with a threshold of *p* < 0.001 uncorrected for multiple comparisons with an extent threshold of 10 voxels were reported. The brain region names were reported with reference to the automated anatomical labeling atlas 3 (AAL3) (Rolls et al., 2020). We additionally compared the averaged parameter estimates (beta values) of ROIs using a two-way within-subject ANOVA with modalities (i.e., visuotactile vs. visuomotor) and timings of visual feedback (i.e., synchronous vs. asynchronous) as factors.

### Multi-voxel pattern analysis (MVPA)

The multivariate classification analysis of fMRI data was performed with a multi-class classifier based on a linear support vector machine implemented in LIBSVM (http://www.csie.ntu.edu.tw/~cjlin/libsvm/) with default parameters (a fixed regularization parameter C = 1). Multi-class classification, implemented in LIBSVM, was used to classify the representations of synchronous and asynchronous conditions. Parameter estimates (beta values) of each trial of voxels within ROIs were used as inputs to the classifier.

We performed two within-modality and one cross-modality classification analyses. Firstly, we ran the within-modality classification analyses between two visuotactile sessions and between two visuomotor sessions. The classifier was first trained to discriminate the representations of the synchronous and asynchronous conditions in the first visuotactile session. The same decoder was then tested to assess whether it could classify the representations of the synchronous and asynchronous conditions in the second visuotactile session. We also conducted a classification in the reverse direction: trained in the second visuotactile session and tested in the first visuotactile session. The averaged decoding accuracy was estimated. The same processing procedure was applied to the within-visuomotor classification analysis. Secondly, we performed a cross-modality classification analysis between the visuotactile and visuomotor sessions. The classifier was first trained to discriminate between the synchronous and asynchronous conditions in the visuotactile sessions. The same decoder was then tested to determine whether it could classify the representations of the synchronous and asynchronous conditions in the visuomotor sessions. We also conducted a classification in the reverse direction: trained in the visuomotor sessions and tested in the visuotactile sessions. The averaged decoding accuracy was estimated. Such a cross-conditional MVPA or cross-classification, a cross-validation between trials with different sets of tasks or stimuli, has been previously used to investigate the similarity or invariance of neural representations by testing the generalization of a classifier between different conditions or modalities (Kaplan et al., 2015). A one-sample *t*-test was used to determine whether the observed decoding accuracy was significantly higher than chance (50%) with intersubject difference treated as a random factor (degrees of freedom = 24). For a ROI-based MVPA, we applied the Holm–Bonferroni procedure (Holm, 1979) based on the number of ROIs in the left and right hemispheres, respectively to control for the problem of multiple comparisons.

Complementary to the a priori ROI analysis, we additionally conducted a volume-based “searchlight” analysis (Kriegeskorte et al., 2006). The classification was performed using multi-voxel activation patterns within a 9-mm-radius sphere (searchlight) that contained 123 voxels. The searchlight moved over the gray matter of the whole brain. The average classification accuracy for each searchlight with cross-validation was assigned to the sphere’s center voxel. The resulting map of the decoding accuracy was averaged over the participants. We used an uncorrected threshold of *p* < 0.001 at the voxel-level and a threshold of *p* < 0.05 FWE corrected at the cluster-level for each type of classification analysis. The names of the brain regions are reported with reference to the AAL3 (Rolls et al., 2020).

## Results

### Questionnaire subjective ratings

The behavioral ratings showed a stronger sense of body ownership in the synchronous condition than in the asynchronous condition. The mean ratings for items (1)–(4) were above 0 in the visuotactile and visuomotor synchronous conditions (visuotactile synchronous condition: [1] 1.16, SD ± 1.64; [2] 1.20, SD ± 2.09; [3] 1.16, SD ± 2.17; [4] 0.68, SD ± 1.92 and visuomotor synchronous condition: [1] 0.52, SD ± 1.97; [2] 0.76, SD ± 2.14; [3] 0.64, SD ± 2.09; [4] 0.04, SD ± 1.76), demonstrating that the participants felt as if the hand on the screen was their own in the synchronous condition. In contrast, the mean ratings for items (5) and (6) were below 0 in the visuotactile and visuomotor asynchronous conditions (visuotactile asynchronous condition: [5] −1.40, SD ± 1.74; [6] −2.04, SD ± 1.18 and visuomotor asynchronous condition: [5] −1.64, SD ± 1.81; [6] −1.24, SD ± 2.10), indicating that the participant’s feeling of ownership for the hand on the screen was less strong than that in the synchronous condition.

Next, the ratings for items (1)–(4) for each condition were averaged to obtain an ownership score and were analyzed by a two-way within-subject ANOVA with modalities (i.e., visuotactile vs. visuomotor) and timings of visual feedback (i.e., synchronous vs. asynchronous) as factors. We found a significant main effect of modalities (*F*(1, 24) = 7.22, *p* = 0.01, *η^2^_p_* = 0.23). The mean ratings were higher in the visuotactile condition than those in the visuomotor condition. We also observed a significant main effect of timings of visual feedback (*F*(1, 24) = 51.12, *p* < 0.01, *η^2^_p_* = 0.68), and the mean ratings were higher in the synchronous condition than they were in the asynchronous condition. There was no significant interaction effect between modalities and timings of visual feedback (*F*(1, 24) = 0.72, *p* = 0.40, *η^2^_p_* = 0.03) (Figure 3).

Lastly, we conducted a paired *t*-test to compare the individual questionnaire items between the synchronous and asynchronous conditions. There were statistically significant or statistical trend toward significant differences (visuotactile synchronous condition vs. visuotactile asynchronous condition: [1] *t* = 5.11, *df* = 24, *p* < 0.01; [2] *t* = 6.18, *df* = 24, *p* < 0.01; [3] *t* = 4.11, *df* = 24, *p* < 0.01; [4] *t* = 4.13, *df* = 24, *p* < 0.01; [5] *t* = −2.22, *df* = 24, *p* = 0.04; [6] *t* = −1.74, *df* = 24, *p* = 0.09; visuomotor synchronous condition vs. visuomotor asynchronous condition: [1] *t* = 5.02, *df* = 24, *p* < 0.01; [2] *t* = 6.72, *df* = 24, *p* < 0.01; [3] *t* = 5.77, *df* = 24, *p* < 0.01; [4] *t* = 5.48, *df* = 24, *p* < 0.01; [6] *t* = −3.15, *df* = 24, *p* < 0.01). The ratings were higher in the synchronous condition than those in the asynchronous condition for items (1)–(4).

**Figure 3:**
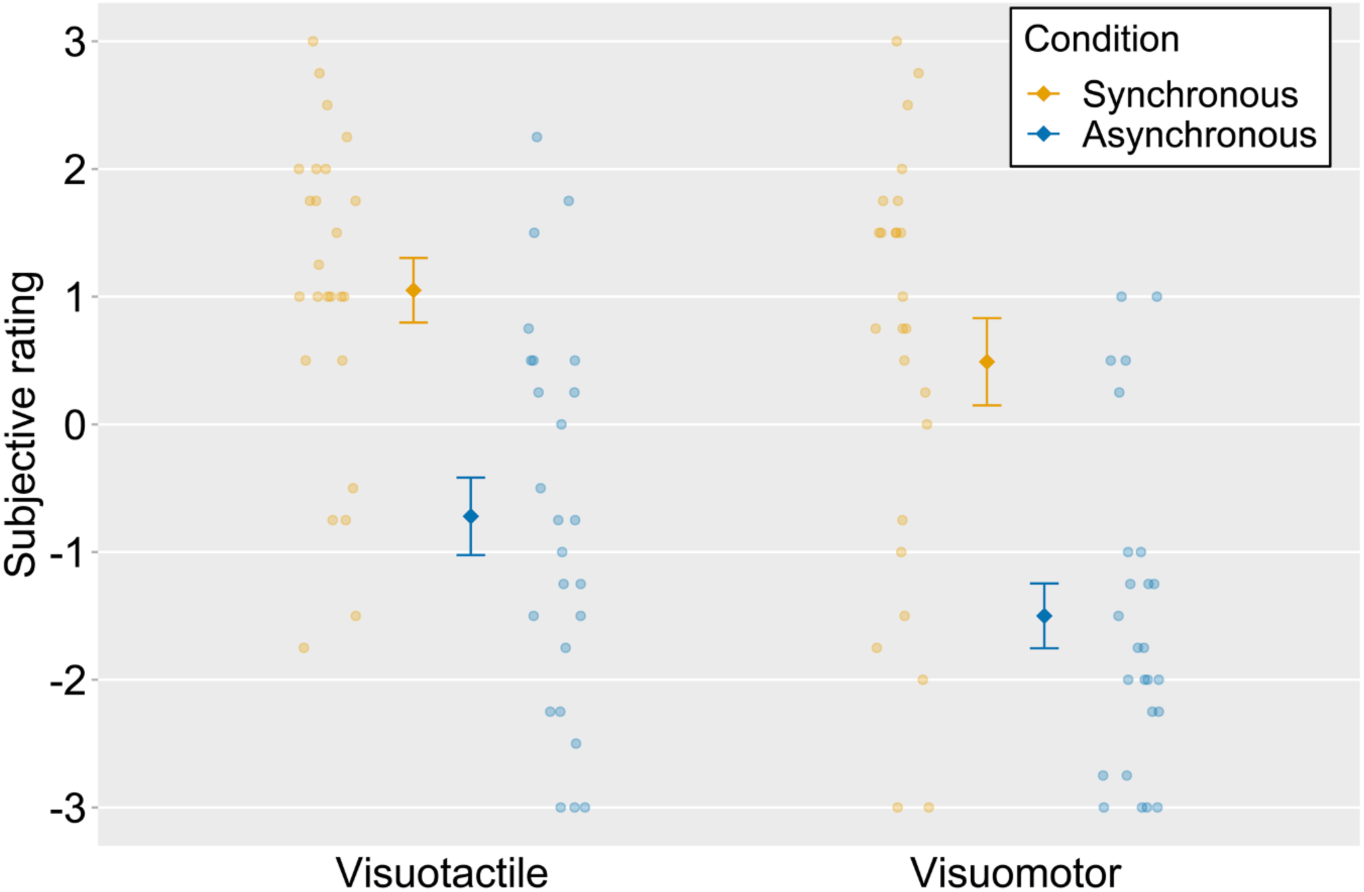
Mean ratings for Ownership questions in four conditions (ownership score). The light-colored circles correspond to the raw data of each participant (*n* = 25) and the means are represented as dark-colored markers. Error bars indicate the standard error of the mean (SEM).

### Mass-univariate analysis

We first compared the activities in the visuotactile and visuomotor conditions collapsed across timings of visual feedback. The activations in the left inferior temporal gyrus, the bilateral postcentral gyrus, and the bilateral rolandic operculum were significantly greater in the visuotactile condition than those in the visuomotor condition. In contrast, the left central sulcus was more activated in the visuomotor condition than in the visuotactile condition. Next, we compared the activities in the synchronous and asynchronous conditions collapsed across modalities. We observed greater activations in the left hippocampus and the right thalamus in the synchronous condition than those in the asynchronous condition. The right triangular part of the inferior frontal gyrus, the left PMv, and the right middle temporal gyrus (MTG) were more activated in the asynchronous condition than in the synchronous condition. The anatomical locations of the activated regions are reported in Table 2 and Table 3.

**Table 2:** Anatomical regions, peak voxel coordinates, and *t*-values of observed activation for the main effect of the visuotactile and visuomotor conditions

| Anatomic region | voxels | MNI coordinates |  |  | *t*-value |
| --- | --- | --- | --- | --- | --- |
|  |  | x | y | z |  |
| *Visuotactile > Visuomotor* |  |  |  |  |  |
| L Inferior temporal gyrus | 101 | −48 | −67 | −7 | 11.94 |
| L Middle occipital gyrus |  | −45 | −70 | 2 | 8.66 |
| L Middle occipital gyrus |  | −39 | −79 | 8 | 6.55 |
| L Postcentral gyrus | 226 | −54 | −16 | 38 | 11.20 |
| L Postcentral gyrus |  | −27 | −37 | 59 | 10.20 |
| L Postcentral sulcus |  | −45 | −28 | 47 | 9.88 |
| R Rolandic operculum | 57 | 45 | −19 | 17 | 11.19 |
| L Rolandic operculum | 130 | −45 | −19 | 20 | 10.99 |
| L Heschl's gyrus |  | −39 | −16 | 11 | 9.43 |
| L Insula |  | −39 | −10 | 5 | 9.31 |
| R Postcentral gyrus | 23 | 57 | −10 | 35 | 8.47 |
| R Postcentral gyrus |  | 60 | −13 | 44 | 7.51 |
| R Postcentral gyrus |  | 51 | −16 | 38 | 6.39 |
| R Postcentral gyrus | 13 | 45 | −25 | 50 | 8.23 |
| R Postcentral gyrus |  | 45 | −22 | 41 | 6.87 |
| L Superior temporal gyrus | 10 | −60 | −4 | 8 | 7.69 |
| L Superior occipital gyrus | 18 | −24 | −76 | 29 | 7.02 |
| L Superior occipital gyrus |  | −21 | −85 | 23 | 6.90 |
| *Visuomotor > Visuotactile* |  |  |  |  |  |
| L Central sulcus | 32 | −33 | −19 | 50 | 8.90 |
| L Central sulcus |  | −30 | −25 | 59 | 7.14 |

Activation was reported with a threshold of $p < 0.05$ corrected for family-wise error (FWE) with an extent threshold of 10 voxels. *MNI*, Montreal Neurological Institute; *L*, left hemisphere; *R*, right hemisphere.

**Table 3:**
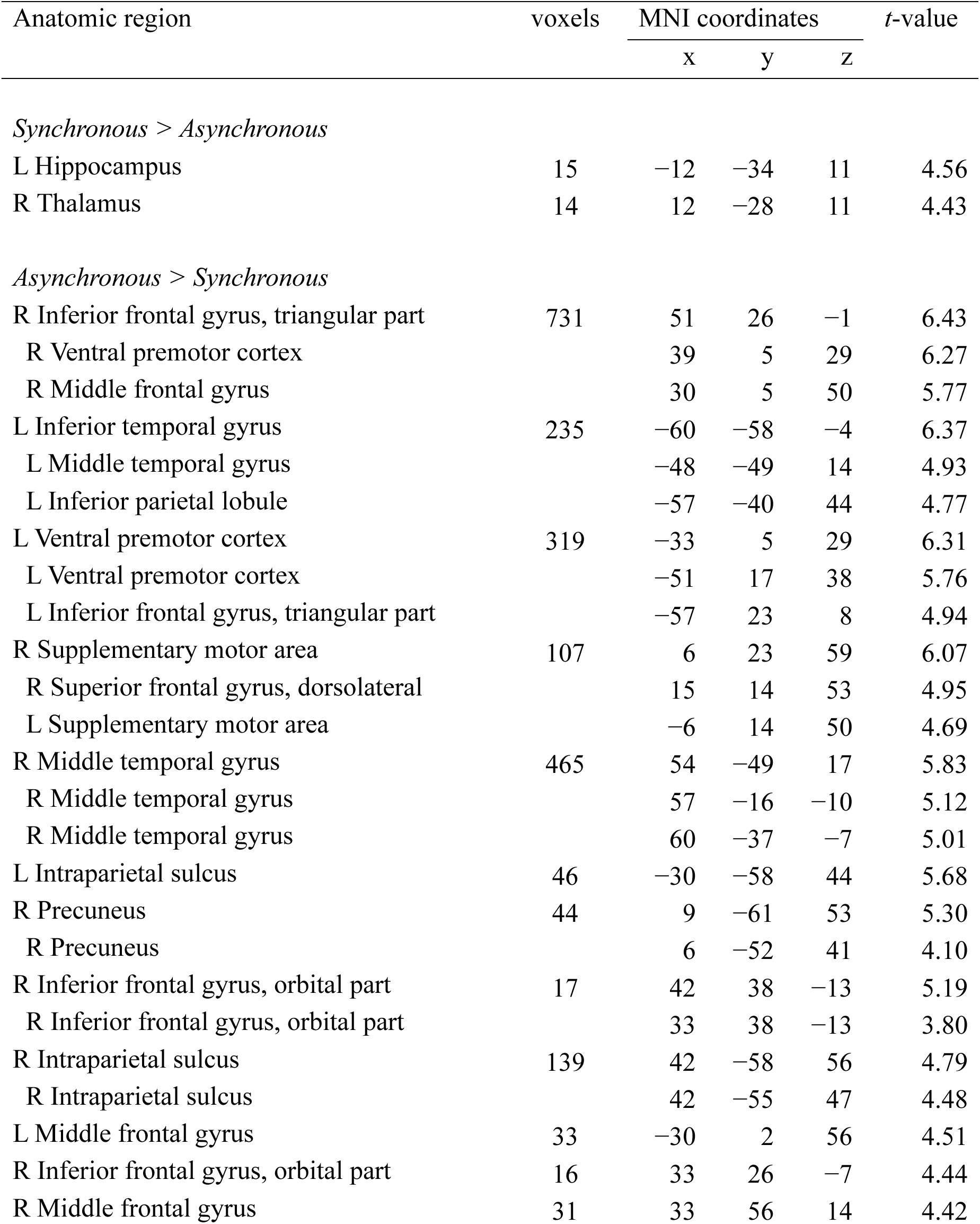
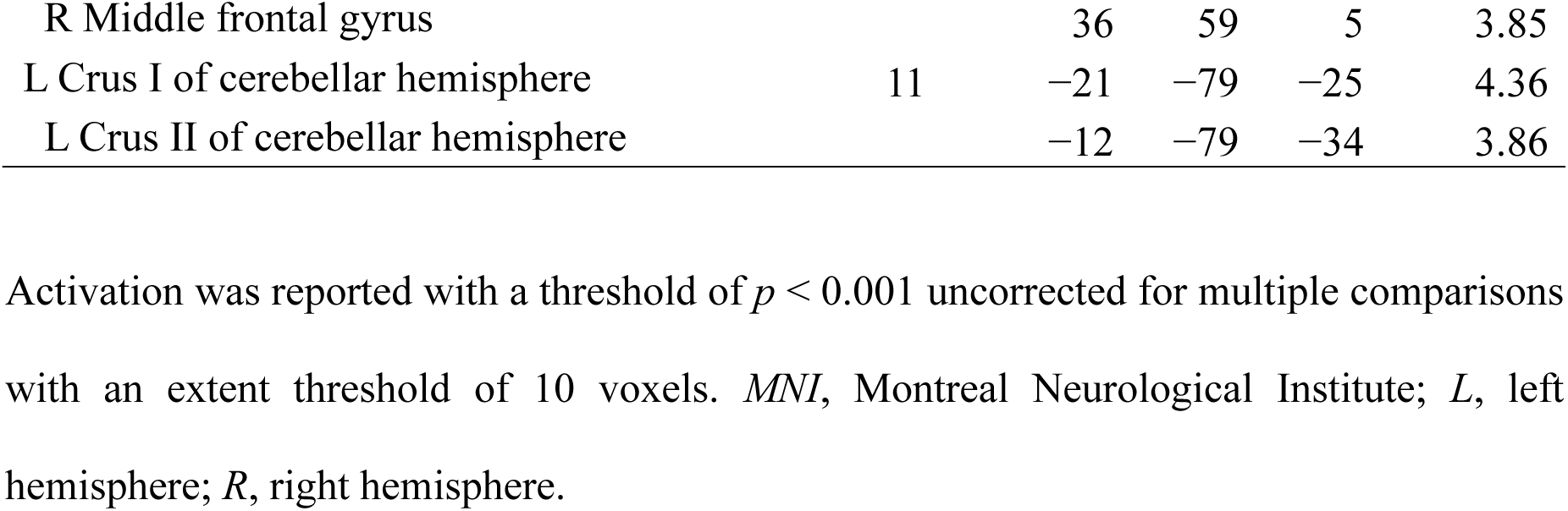
Anatomical regions, peak voxel coordinates, and *t*-values of observed activation for the main effect of synchronous and asynchronous conditions

| Anatomic region | voxels | MNI coordinates |  |  | *t*-value |
| --- | --- | --- | --- | --- | --- |
|  |  | x | y | z |  |
| *Synchronous > Asynchronous* |  |  |  |  |  |
| L Hippocampus | 15 | −12 | −34 | 11 | 4.56 |
| R Thalamus | 14 | 12 | −28 | 11 | 4.43 |
| *Asynchronous > Synchronous* |  |  |  |  |  |
| R Inferior frontal gyrus, triangular part | 731 | 51 | 26 | −1 | 6.43 |
| R Ventral premotor cortex |  | 39 | 5 | 29 | 6.27 |
| R Middle frontal gyrus |  | 30 | 5 | 50 | 5.77 |
| L Inferior temporal gyrus | 235 | −60 | −58 | −4 | 6.37 |
| L Middle temporal gyrus |  | −48 | −49 | 14 | 4.93 |
| L Inferior parietal lobule |  | −57 | −40 | 44 | 4.77 |
| L Ventral premotor cortex | 319 | −33 | 5 | 29 | 6.31 |
| L Ventral premotor cortex |  | −51 | 17 | 38 | 5.76 |
| L Inferior frontal gyrus, triangular part |  | −57 | 23 | 8 | 4.94 |
| R Supplementary motor area | 107 | 6 | 23 | 59 | 6.07 |
| R Superior frontal gyrus, dorsolateral |  | 15 | 14 | 53 | 4.95 |
| L Supplementary motor area |  | −6 | 14 | 50 | 4.69 |
| R Middle temporal gyrus | 465 | 54 | −49 | 17 | 5.83 |
| R Middle temporal gyrus |  | 57 | −16 | −10 | 5.12 |
| R Middle temporal gyrus |  | 60 | −37 | −7 | 5.01 |
| L Intraparietal sulcus | 46 | −30 | −58 | 44 | 5.68 |
| R Precuneus | 44 | 9 | −61 | 53 | 5.30 |
| R Precuneus |  | 6 | −52 | 41 | 4.10 |
| R Inferior frontal gyrus, orbital part | 17 | 42 | 38 | −13 | 5.19 |
| R Inferior frontal gyrus, orbital part |  | 33 | 38 | −13 | 3.80 |
| R Intraparietal sulcus | 139 | 42 | −58 | 56 | 4.79 |
| R Intraparietal sulcus |  | 42 | −55 | 47 | 4.48 |
| L Middle frontal gyrus | 33 | −30 | 2 | 56 | 4.51 |
| R Inferior frontal gyrus, orbital part | 16 | 33 | 26 | −7 | 4.44 |
| R Middle frontal gyrus | 31 | 33 | 56 | 14 | 4.42 |
| R Middle frontal gyrus |  | 36 | 59 | 5 | 3.85 |
| L Crus I of cerebellar hemisphere | 11 | -21 | -79 | -25 | 4.36 |
| L Crus II of cerebellar hemisphere |  | -12 | -79 | -34 | 3.86 |

Activation was reported with a threshold of $p < 0.001$ uncorrected for multiple comparisons with an extent threshold of 10 voxels. *MNI*, Montreal Neurological Institute; *L*, left hemisphere; *R*, right hemisphere.

We then investigated the areas showing interaction effects between modalities and timings of visual feedback. In the visuotactile condition, the difference between synchronous and asynchronous conditions was significantly larger in the left precuneus, the bilateral cerebellum, and the right fusiform gyrus than that in the visuomotor condition. In the visuomotor condition, the difference between the synchronous and asynchronous conditions was significantly greater in the left inferior parietal lobule than that in the visuotactile condition. The anatomical locations of the activated areas are reported in Table 4 and Figure 4.

**Figure 4:**
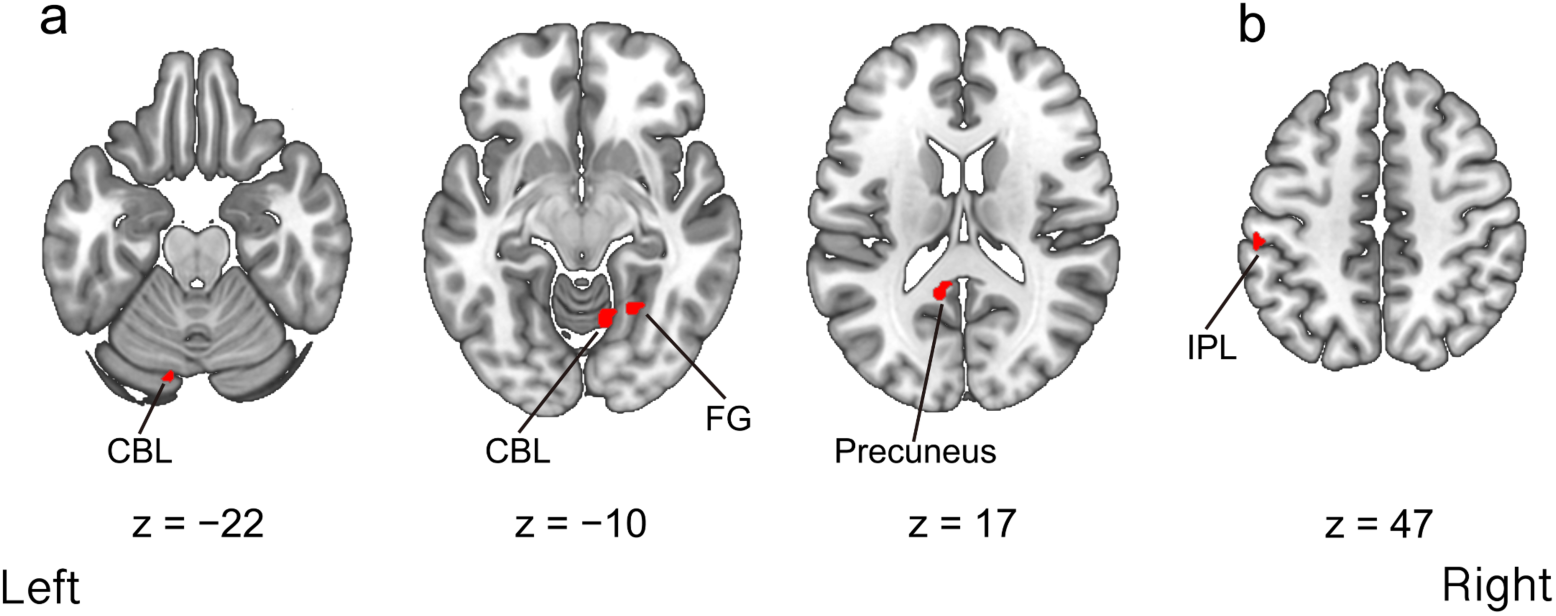
Regions activated by the interactions of modalities and timings of visual feedback in the fMRI univariate analysis. **a**, Activated regions that showed the greater difference between the visuotactile synchronous and asynchronous conditions compared with the difference between the visuomotor synchronous and asynchronous conditions. **b**, Activated regions that showed the greater difference between the visuomotor synchronous and asynchronous conditions compared with the difference between the visuotactile synchronous and asynchronous conditions. Activation was reported with a threshold of *p* < 0.001 uncorrected for multiple comparisons with an extent threshold of 10 voxels. Montreal Neurological Institute (MNI) coordinates of the activated foci are reported in Table 4. CBL, cerebellum; FG, fusiform gyrus; IPL, inferior parietal lobule.

**Table 4:** Anatomical regions, peak voxel coordinates, and *t*-values of observed activation for the interaction effects

| Anatomic region | voxels | MNI coordinates |  |  | *t*-value |
| --- | --- | --- | --- | --- | --- |
|  |  | x | y | z |  |
| *Visuotactile synchronous interaction* |  |  |  |  |  |
| L Precuneus | 12 | −9 | −49 | 17 | 4.70 |
| L Crus I of cerebellar hemisphere | 11 | −12 | −82 | −22 | 4.05 |
| R Lobule VI of cerebellar hemisphere | 16 | 12 | −61 | −10 | 4.03 |
| R Lingual gyrus |  | 18 | −67 | −4 | 3.72 |
| R Fusiform gyrus | 13 | 24 | −55 | −10 | 3.96 |
| *Visuomotor synchronous interaction* |  |  |  |  |  |
| L Inferior parietal lobule | 20 | −57 | −28 | 47 | 4.99 |
| L Inferior parietal lobule |  | −54 | −31 | 38 | 3.82 |

Activation was reported with a threshold of $p < 0.001$ uncorrected for multiple comparisons with an extent threshold of 10 voxels. *MNI*, Montreal Neurological Institute; *L*, left hemisphere; *R*, right hemisphere.

The fMRI data analysis of the EBA localizer scan revealed a significant activation of the left middle occipital gyrus and the right MTG when the participants watched body parts pictures compared to the activity elicited by chair images (MNI coordinates: [−48, −70, 8] for the left hemisphere and [51, −58, 2] for the right hemisphere). The anatomical locations of the activated areas are indicated in Table 5 and Figure 5. We also assessed the number of correct responses, namely pressing the button when the same images appeared twice successively in the body part and chair image blocks, to confirm whether the participants watched the stimulus. The results showed that the percentage of correct responses was 97.6%, indicating that the participants watched the stimulus closely.

**Figure 5:**
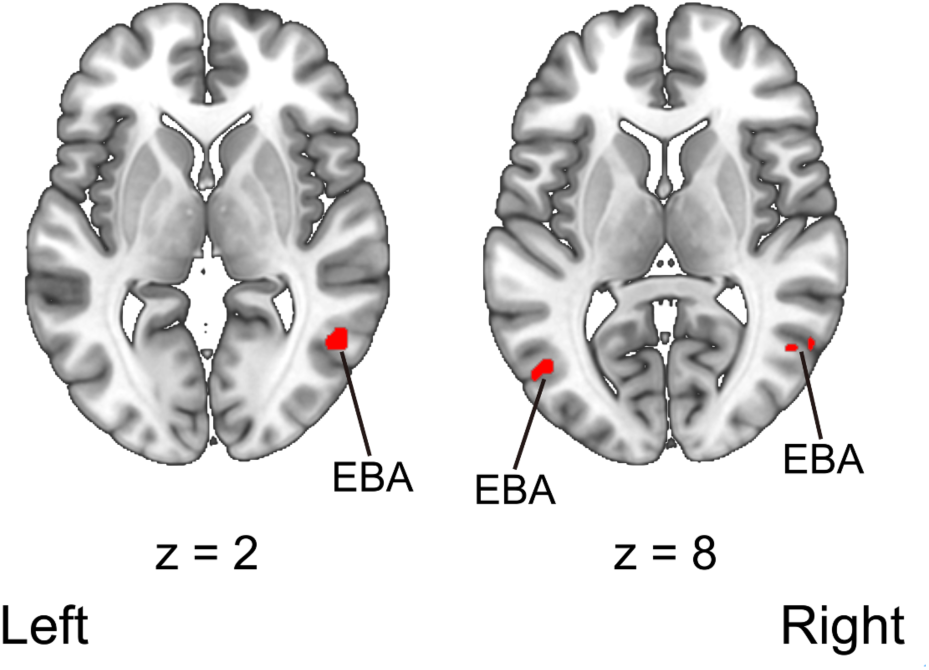
Regions activated by the presentation of body parts images compared with that of chair images during the EBA localizer scan. Activation was reported with a threshold of *p* < 0.05 corrected for family-wise error (FWE) with an extent threshold of 10 voxels. Montreal Neurological Institute (MNI) coordinates of the activated foci are reported in Table 5. EBA, extrastriate body area.

**Table 5:**
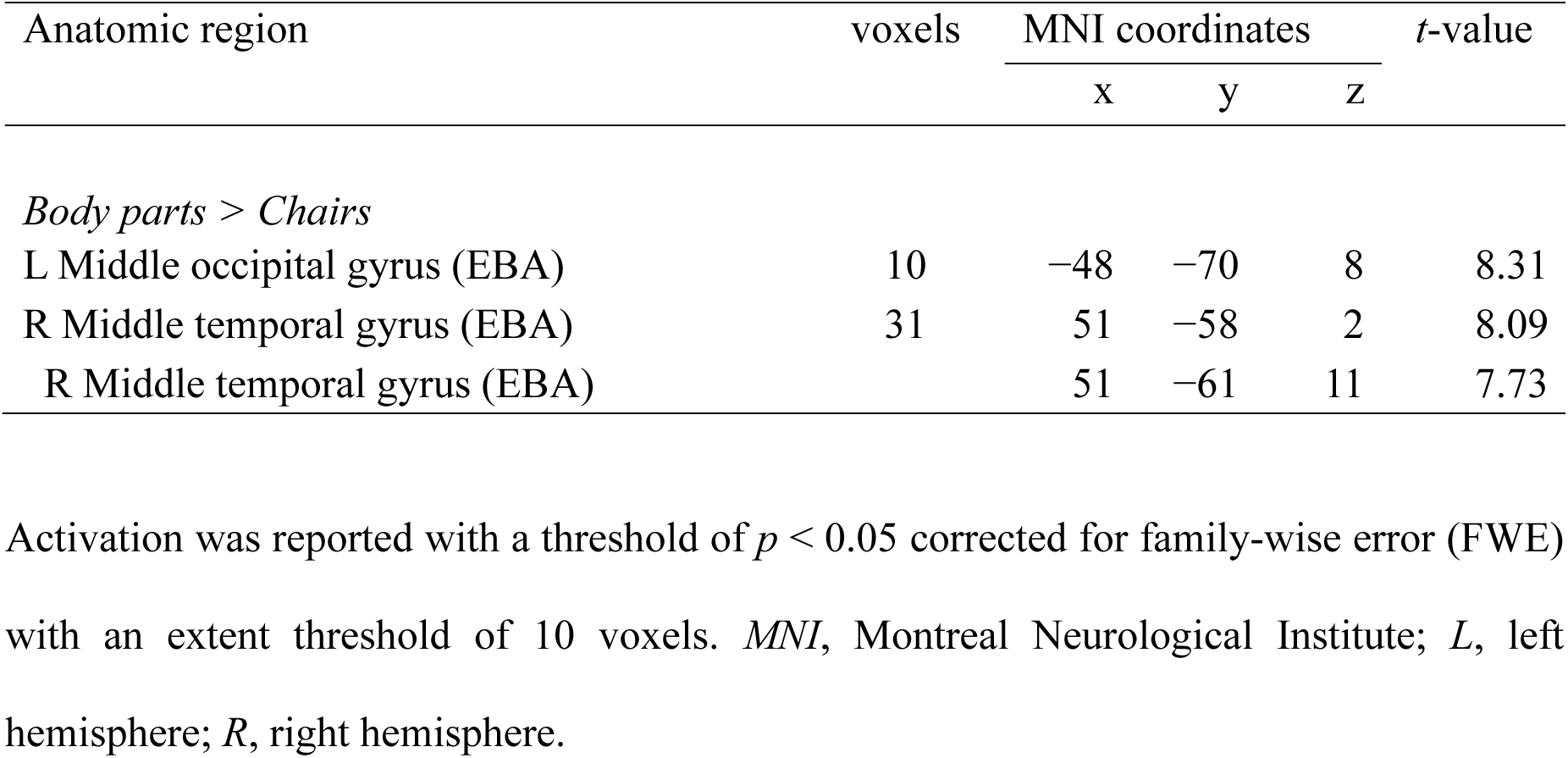
Anatomical regions, peak voxel coordinates, and *t*-values of observed activation during the EBA localizer scan

Next, we extracted the parameter estimates of ROIs for each condition. The averaged parameter estimates (beta values) were analyzed using a two-way within-subject ANOVA with modalities (i.e., visuotactile vs. visuomotor) and timings of visual feedback (i.e., synchronous vs. asynchronous) as factors. There was a significant main effect of the modalities in the bilateral IPS (*F* (1, 24) = 9.32, *p* = 0.01, *η^2^_p_* = 0.28 for the left hemisphere and *F*(1, 24) = 7.47, *p* = 0.01, *η^2^_p_* = 0.24 for the right hemisphere), the bilateral PMv (*F*(1, 24) = 8.77, *p* = 0.01, *η^2^_p_* = 0.27 for the left hemisphere and *F*(1, 24) = 6.19, *p* = 0.02, *η^2^_p_* = 0.21 for the right hemisphere), and the left EBA (*F*(1, 24) = 46.99, *p* < 0.01, *η^2^_p_* = 0.66). The averaged parameter estimates were greater in the visuotactile condition than those in the visuomotor condition.

We also found a significant main effect of the timings of the visual feedback in the bilateral IPS (*F*(1, 24) = 9.65, *p* < 0.01, *η^2^_p_* = 0.29 for the left hemisphere and *F*(1, 24) = 13.65, *p* < 0.01, *η^2^_p_* = 0.36 for the right hemisphere) and the bilateral PMv (*F*(1,24) = 10.16, *p* < 0.01, *η^2^_p_* = 0.30 for the left hemisphere and *F*(1,24) = 8.51, *p* = 0.01, *η^2^_p_* = 0.26 for the right hemisphere). There was a statistical trend toward a main effect of the timings of the visual feedback for the right EBA (*F*(1, 24) = 3.21, *p* = 0.09, *η^2^_p_* = 0.12). The parameter estimates in the aforementioned ROIs were significantly stronger in the asynchronous condition than those in the synchronous condition. No significant interaction between modalities and timings of visual feedback was observed in all ROIs (*Fs*(1, 24) < 0.14, *ps* > 0.71, *η^2^_p_s* ≦ 0.01) (Figure 6).

**Figure 6:**
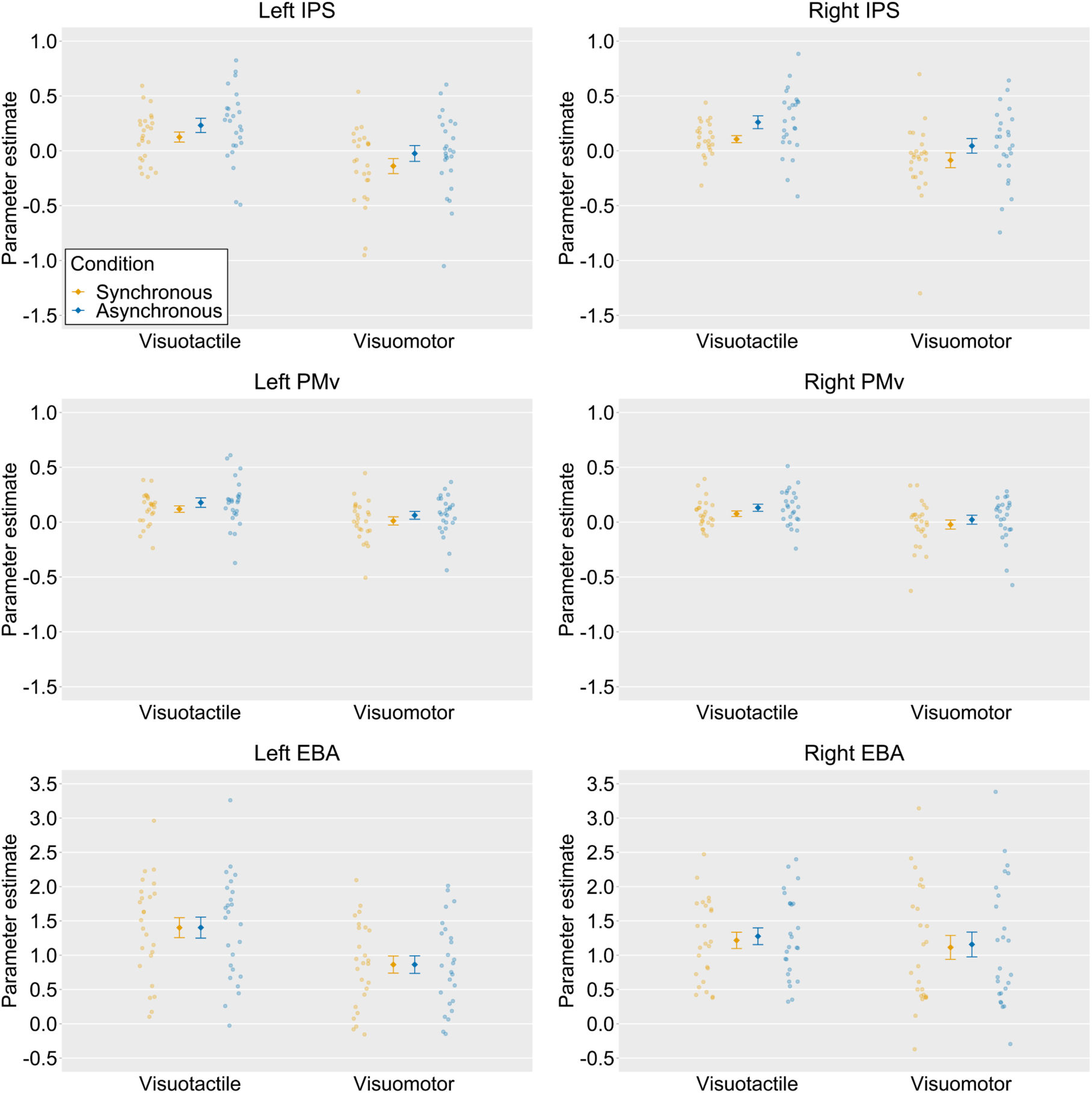
Averaged activation (parameter estimates) within ROIs. The light-colored circles correspond to the raw data of each participant (*n* = 25) and the means are represented as dark-colored markers. Error bars indicate SEM. IPS, intraparietal sulcus; PMv, ventral premotor cortex; EBA, extrastriate body area.

### MVPA

We first conducted a ROI-based MVPA to classify the representations of the synchronous and asynchronous conditions within the same and across modalities. Within the visuotactile classification, a significant above-chance decoding accuracy was found for the right PMv (54.12%, *t*(24) = 2.37, *p* = 0.03, Cohen’s *d* = 0.48) and a trend toward statistical significance was obtained for the left PMv (53.88%, *t*(24) = 1.98, *p* = 0.06, Cohen’s *d* = 0.40). Both these results were not significant after threshold correction for multiple comparisons. Within the visuomotor classification, the classification accuracy was significant for the left IPS (55.00%, *t*(24) = 2.79, *p* = 0.01, Cohen’s *d* = 0.57) and the left EBA (55.12%, *t*(24) = 2.98, *p* = 0.01, Cohen’s *d* = 0.61). There was a trend toward statistical significance for the decoding accuracy in the right PMv (53.12%, *t*(24) = 2.00, *p* = 0.06, Cohen’s *d* = 0.41). Among these ROIs, the decoding accuracies in the left IPS and the left EBA were significant after correction for multiple comparisons. In cross-classification between visuotactile and visuomotor sessions, significant above-chance decoding accuracies were found for the bilateral IPS (53.56%, *t*(24) = 2.77, *p* = 0.01, Cohen’s *d* = 0.57 for the left hemisphere and 54.19%, *t*(24) = 3.54, *p* < 0.01, Cohen’s *d* = 0.72 for the right hemisphere) and the left PMv (53.06%, *t*(24) = 2.83, *p* = 0.01, Cohen’s *d* = 0.58), and the three ROIs showed significant above-chance accuracy after correction for multiple comparisons (Figure 7).

**Figure 7:**
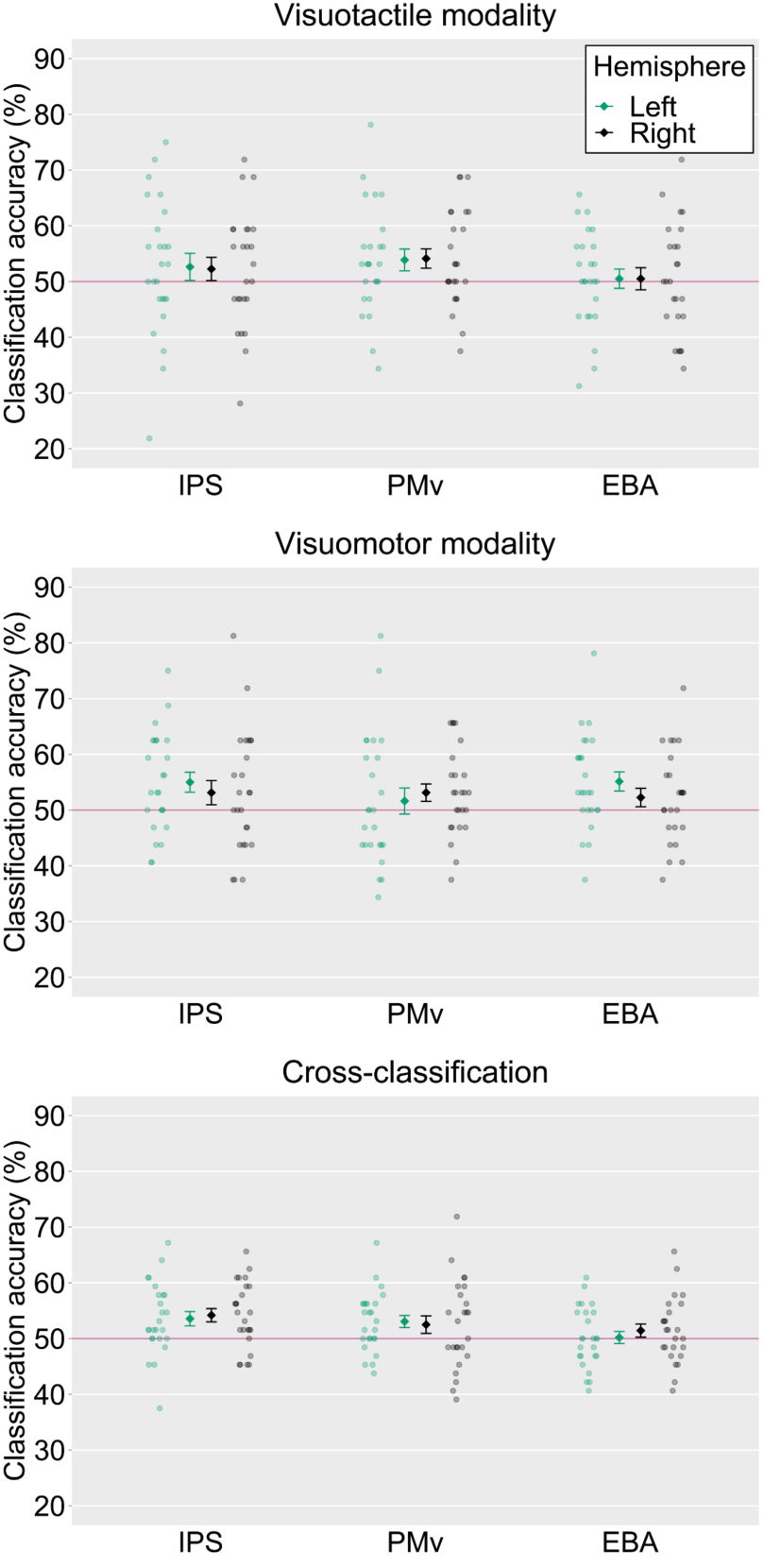
Averaged classification accuracies in each ROI. The light-colored circles correspond to the raw data of each participant (*n* = 25) and the means are represented as dark-colored markers. Error bars indicate SEM. Horizontal red lines represent chance-level (50%). IPS, intraparietal sulcus; PMv, ventral premotor cortex; EBA, extrastriate body area.

Additionally, a voxel bias map was generated to investigate whether the neural patterns decoded by MVPA were similar among participants. The voxel bias map displays the average weights (positive or negative) of the classifier across two folds (i.e., iterations) in the cross-validation for individual voxels within ROIs. This map showed intermingled patterns of voxels with two timings of visual feedback (synchronous vs. asynchronous biased voxels) within ROIs in all subjects (Supplementary Figure 1A). In addition, the bias patterns between different pairs of participants revealed low correlations around zero (Supplementary Figure 1B), indicating idiosyncratic patterns of weights specific to each subject.

We then ran searchlight analyses to decode the timings of the visual feedback (i.e., synchronous vs. asynchronous) within the same and across modalities. Firstly, significantly above-chance classification accuracies were found in the right IPS, the right PMv, and the right postcentral sulcus within visuotactile sessions. Secondly, we also observed a significant above-chance classification accuracy in the left central sulcus within-visuomotor sessions. Thirdly, the analysis revealed significant above-chance classification accuracies in the right PMv, the left EBA, and the bilateral IPS in cross-classification between visuotactile and visuomotor sessions. The anatomical locations of the activated areas are reported in Table 6 and Figure 8.

**Figure 8:**
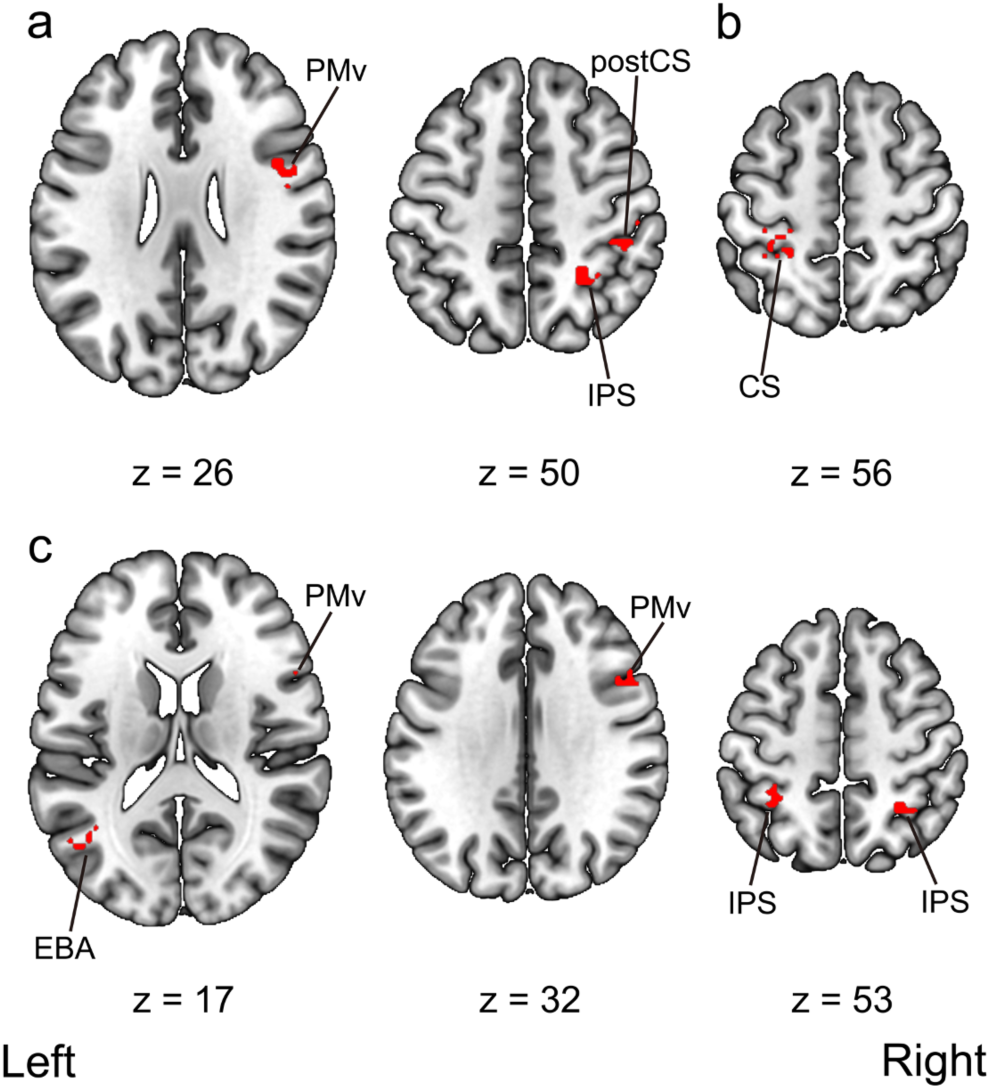
Multi-voxel pattern analysis searchlight results. **a**, Regions that showed significant above-chance decoding accuracy within the visuotactile classification. **b**, Regions that showed significant above-chance decoding accuracy within the visuomotor classification. **c**, Regions that showed significant above-chance cross-classification accuracy between the visuotactile and visuomotor classification. Activation was reported with an uncorrected threshold of *p* < 0.001 at the voxel-level and a threshold of *p* < 0.05 family-wise error (FWE) corrected at the cluster-level. Montreal Neurological Institute (MNI) coordinates of the activated foci are reported in Table 6. PMv, ventral premotor cortex; IPS, intraparietal sulcus; postCS, postcentral sulcus; CS, central sulcus; EBA, extrastriate body area. The unthresholded raw *t*-value maps in NIfTY format are available at https://doi.org/10.6084/m9.figshare.19228050.

**Table 6:**
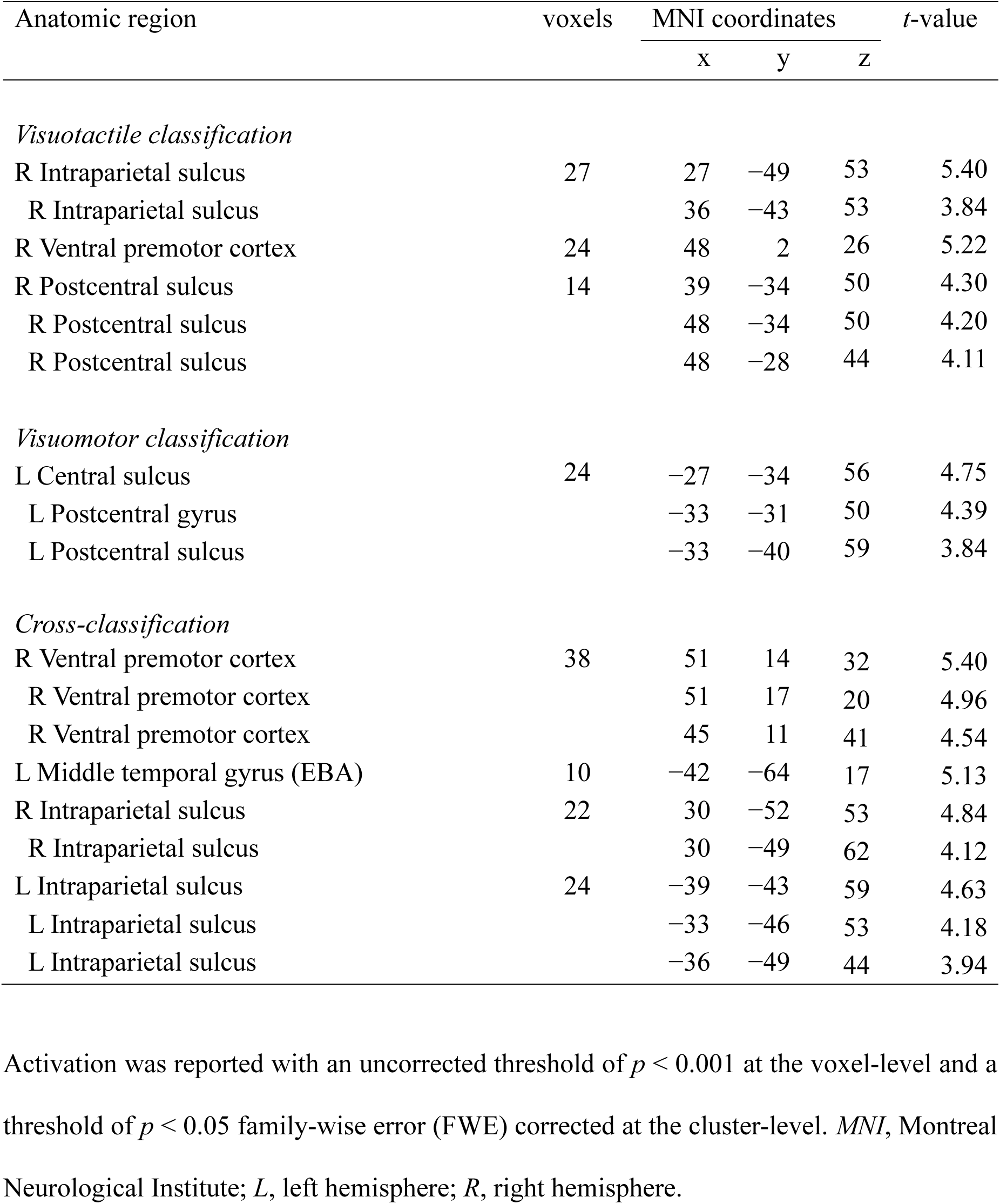
Anatomical regions, peak voxel coordinates, and *t*-values of observed activation of the searchlight multi-voxel pattern analysis

### Correlations between cross-classification accuracy and subjective rating

We analyzed the relationship between the subjective ratings in the questionnaire and the cross-classification accuracies. We first averaged the responses to items (1)–(4) in the visuotactile synchronous and visuomotor synchronous condition and in the visuotactile asynchronous and visuomotor asynchronous condition. We then calculated the differences between the average scores obtained in the synchronous condition and those obtained in the asynchronous condition and correlated the values of these differences with the cross-classification accuracies in each ROI. We found a significant positive correlation between the value and the cross-classification accuracy for the left PMv (Pearson’s *r* = 0.54, *n* = 25, *p* = 0.01) and a trend toward statistical significance for the right IPS (Pearson’s *r* = 0.38, *n* = 25, *p* = 0.06). A significant positive correlation was observed for the left PMv after correction for multiple comparisons (Figure 9).

**Figure 9:**
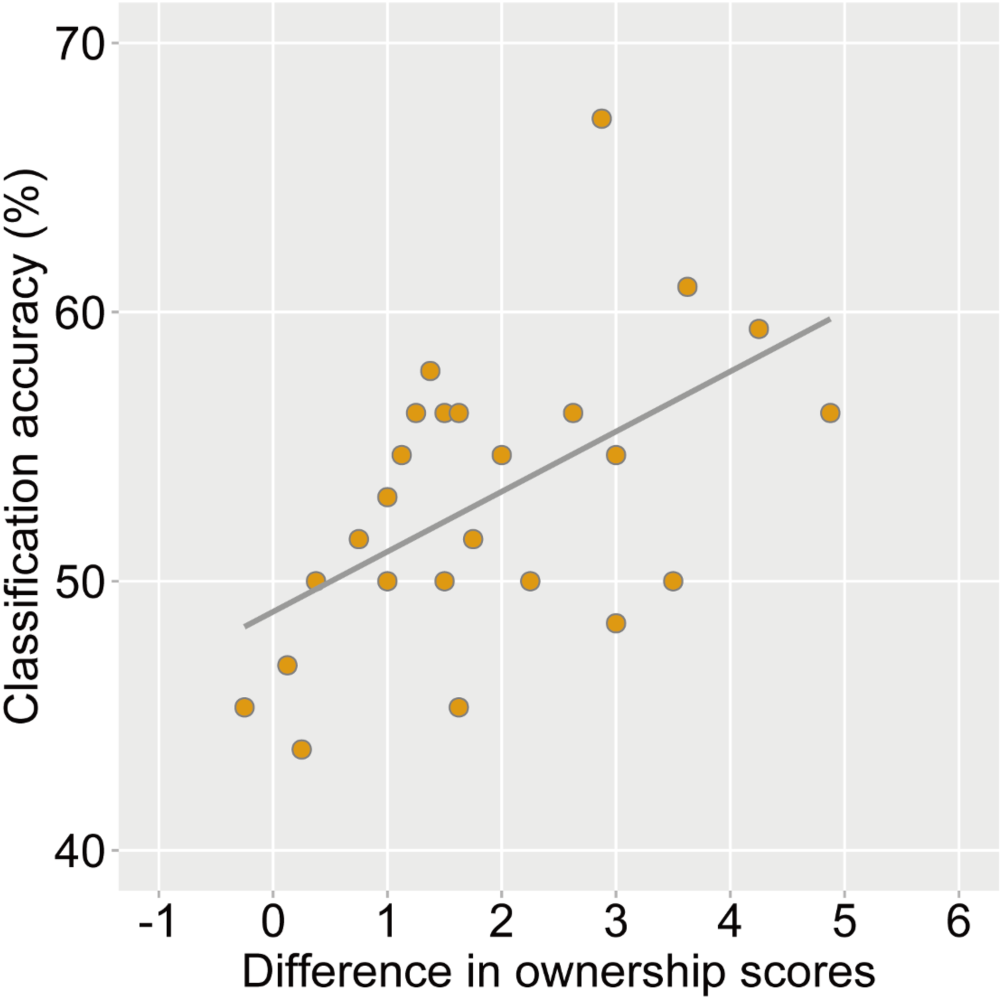
Correlation analysis between the cross-classification accuracy in the left ventral premotor cortex (PMv) and the difference in subjective ratings between the synchronous and asynchronous conditions (Pearson’s *r* = 0.54, *n* = 25, *p* = 0.01). Each circle corresponds to the data of each participant (*n* = 25).

## Discussion

The present study tried to reveal the supramodal neural representations of the sense of body ownership for visuotactile and visuomotor modalities. We manipulated the sense of body ownership using visuotactile and visuomotor inputs within a single fMRI experiment and assessed whether the classifier first trained on the data from the visuotactile modality could decode the timings of the visual feedback (i.e., synchronous vs. asynchronous) on the data from the visuomotor modality and vice versa. This cross-classification analysis revealed that the IPS, PMv, and EBA subserve the neural representations of the sense of body ownership common to the visuotactile and visuomotor modalities. We also found a statistically significant correlation between the cross-classification accuracy in the left PMv and the difference in subjective ratings between the synchronous and asynchronous conditions. These findings indicate that the sense of body ownership is represented regardless of the modalities in the IPS, PMv, and EBA.

The IPS and PMv exhibited significantly higher cross-classification accuracies for visuotactile and visuomotor modalities in ROI-based MVPA. Such high decoding accuracies were further obtained by searchlight MVPA, providing compelling evidence that there are neural patterns of the sense of body ownership invariant to the modalities in the parieto-premotor cortices. Previous studies showed the role of the IPS and PMv in multisensory integration and the sense of body ownership (Ehrsson et al., 2004; Gentile et al., 2015; Limanowski and Blankenburg, 2015, 2016a, 2018; Grivaz et al., 2017). The role of these regions in the sense of body ownership was also suggested by a lesion study (Zeller et al., 2011) and studies using transcranial magnetic stimulation and transcranial direct current stimulation of these regions (Karabanov et al., 2017; Convento et al., 2018; Lira et al., 2018). Consistent with previous findings, our study highlights the neural networks involved in the abstract sense of body ownership induced by multisensory information.

Our searchlight analysis revealed above-chance cross-classification accuracy for the left EBA. The EBA is a region of the LOC involved in the visual processing of the human body such as the perception of one’s movement (Astafiev et al., 2004) and mental imagery of the human body (Blanke et al., 2010). Previous neuroimaging studies in humans indicated an increased activation in the LOC, especially in the EBA, to the sense of body ownership (Limanowski et al., 2014; Wold et al., 2014; Limanowski and Blankenburg, 2015, 2016a). However, compared with the IPS and PMv, which were the focus of electrophysiological studies in macaques and neuroimaging studies in humans (Graziano et al., 1997; Duhamel et al., 1998; Ehrsson et al., 2004; Avillac et al., 2007; Tsakiris et al., 2010; Gentile et al., 2011; Olivé et al., 2015), the role of the EBA in the sense of body ownership has not been fully clarified. Our results provide further insight and reveal a supramodal signature of the sense of body ownership across the sensory modalities in the EBA. However, no significant cross-classification accuracy was observed for the EBA in the ROI-based MVPA, which was performed with ROIs defined with the localizer session. This result inconsistency might be due to the difference in the voxels included in ROIs in the ROI-based and searchlight analyses. Altogether, the data suggest that the supramodal sense of body ownership is represented in the IPS, PMv, and EBA.

The univariate analysis revealed that the averaged activities in the bilateral IPS and PMv were significantly greater in the asynchronous condition than those in the synchronous condition. This seems in disagreement with data from previous RHI studies (Ehrsson et al., 2004; Limanowski and Blankenburg, 2015). However, this discrepancy is likely explained by the differences in the experimental settings. Previous studies (Ehrsson et al., 2004; Limanowski and Blankenburg, 2015) used a fake hand instead of a real one. Thus, visual stimuli might not elicit the sense of body ownership by default in participants and experimental manipulation might be required for participants to feel body ownership. In contrast, in the present study, the participants watched their own video-recorded hand. This might trigger the sense of ownership for the hand displayed on the screen. Interestingly, the results of the present study are consistent with those obtained by Tsakiris et al. (2010) using a similar procedure. Furthermore, Gentile et al. (2013), who recorded videos of tactile stimuli applied to the participants’ own hands before the MRI scan and used them as visual stimuli during the MRI scan, reported that averaged activities in the PMv, IPS, and LOC were greater in the synchronous condition like the previous studies using a fake hand (e.g., Limanowski and Blankenburg, 2015). Therefore, it is possible that regions such as the IPS, PMv, and EBA represent the extent to which the “default” sense of body ownership for visual stimuli has been updated by experimental manipulations (i.e., synchronous or asynchronous stimulation). The use of a fake hand or recordings of a real hand (Ehrsson et al., 2004; Gentile et al., 2013; Limanowski and Blankenburg, 2015) may result in larger updates in the synchronous condition, while the presentation of a real hand in real-time (Tsakiris et al., 2010) larger updates in the asynchronous condition. A theoretical model explaining RHI from a predictive coding framework (Apps and Tsakiris, 2014) also suggested that these regions are associated with the default state (in the model, called “empirical prior”) of body ownership. This interpretation is consistent with the claim that the sense of body ownership over the fake hand and the (dis)ownership of the real hand have a common neural basis (Ehrsson, 2020).

Although our results revealed the existence of a shared neural representation of the sense of body ownership across the modalities, our behavioral ratings showed that the sense of body ownership was elicited more strongly in the visuotactile condition than it was in the visuomotor condition. This might be caused by the difference in the threshold detection of the visual feedback delay between the visuotactile and visuomotor conditions. In the present study, the sense of body ownership in the visuotactile condition was induced by visual and proprioceptive information, whereas the sense of body ownership in the visuomotor condition was caused by various afferent signals arising from the muscle spindles, tendon organ, and joint receptors (Proske and Gandevia, 2012). Thus, the visuomotor condition elicited not only the sense of body ownership but also the sense of agency that was not induced in the visuotactile condition (Kalckert and Ehrsson, 2014b). Therefore, in the visuomotor condition, the perception of the mismatch between the intention to move the finger and the visual feedback allowed the participants to detect the visual feedback delay and to determine whether the hand on the screen was their own.

Another possible interpretation of the difference in the behavioral ratings comes from the constraints on the experimental design. While we executed enough trials for MVPA, the stimulus presentation was shorter (i.e., 18 s) than that in previous studies (Ehrsson et al., 2004, 2007; Chae et al., 2015; Olivé et al., 2015; Lee and Chae, 2016). Recent behavioral evidence suggested that the ownership sensation for visuomotor RHI takes approximately 23 s to emerge (Kalckert and Ehrsson, 2017). Thus, the duration of the stimulus presentation might have been insufficient for the ownership sensation to emerge, especially in the visuomotor condition, leading to a smaller ownership score in the visuomotor condition than that in the visuotactile condition. Future studies are needed to establish whether the sense of body ownership is induced to the same degree and to determine the stimulation duration necessary to elicit the ownership sensation under different conditions.

The current study also showed the lower subjective ratings in the synchronous and asynchronous conditions compared to the previous fMRI study with similar experimental settings (Tsakiris et al., 2010). The result indicates that the participants might not have felt a strong sense of body ownership, even with a visual feedback of their real hand. This might be due to the intrinsic delay that was inevitable when the videos captured by the camera were presented in real-time. Temporal discrepancies between the tactile stimulation of the real hand or finger movements and the visual feedback caused multisensory integration to break down leading to disembodiment (Roel Lesur et al., 2020). The intrinsic delay of approximately 400 ms might have hindered the participants from feeling the hand on the screen as their own, even in the synchronous condition. This can be considered as a limitation to the present study. However, two-way within-subject ANOVA revealed a significant difference in the subjective ratings between the synchronous and asynchronous conditions. We also found higher ratings in the synchronous condition for individual questionnaire items (1)–(4). Thus, the participants perceived the contrast of the visual feedback between the synchronous and asynchronous conditions and judged whether the hand on the screen was their own based on the temporal (in)congruency among sensory signals.

The current study has another methodological limitation. Indeed, in the visuomotor condition, the participants received tactile stimulation of the palmar side of the index finger when they executed finger movements. Therefore, a tactile stimulation was present in the visuomotor and the visuotactile conditions. Although this might result in the successful cross-classification between the conditions, we considered that it was unlikely for the following reasons. Firstly, the stimulated body part was different between the conditions: the tactile stimulation in the visuotactile condition consisted in stroking the back of the finger with a paintbrush, while that in the visuomotor condition was on the palmer side of the finger. In addition, as self-generated movements have been reported to attenuate sensory feedback (e.g., Blakemore et al., 1998), the stimulation in the visuomotor condition was perceived as weaker than that in the visuotactile condition. Therefore, the successful cross-classification cannot be explained only by the tactile inputs and indicates the sense of body ownership regardless of the modalities.

In conclusion, the present study investigated the supramodal neural representations of the sense of body ownership induced by different combinations of sensory inputs. We demonstrated a shared neural representation of the sense of body ownership in the IPS, PMv, EBA for visuotactile and visuomotor modalities. Furthermore, we revealed that the cross-classification accuracy in the left PMv significantly positively correlated with the difference in subjective ratings of the sense of body ownership between synchronous and asynchronous conditions. Our findings provide novel insights into the integration of bodily signals mediating the sense of body ownership.

## Acknowledgments

This work was supported by JSPS KAKENHI Grant Numbers 18H01098, 19H00634 & 21H00958 to K.O. and partially supported by Graduate Grant Program of Graduate School of Humanities and Human Sciences, Hokkaido University, awarded to T.Y. The authors would like to thank Enago (www.enago.jp) for the English language review.

## Conflict of interest

The authors declare no competing financial interests.

---

**Supplementary Figure 1:**
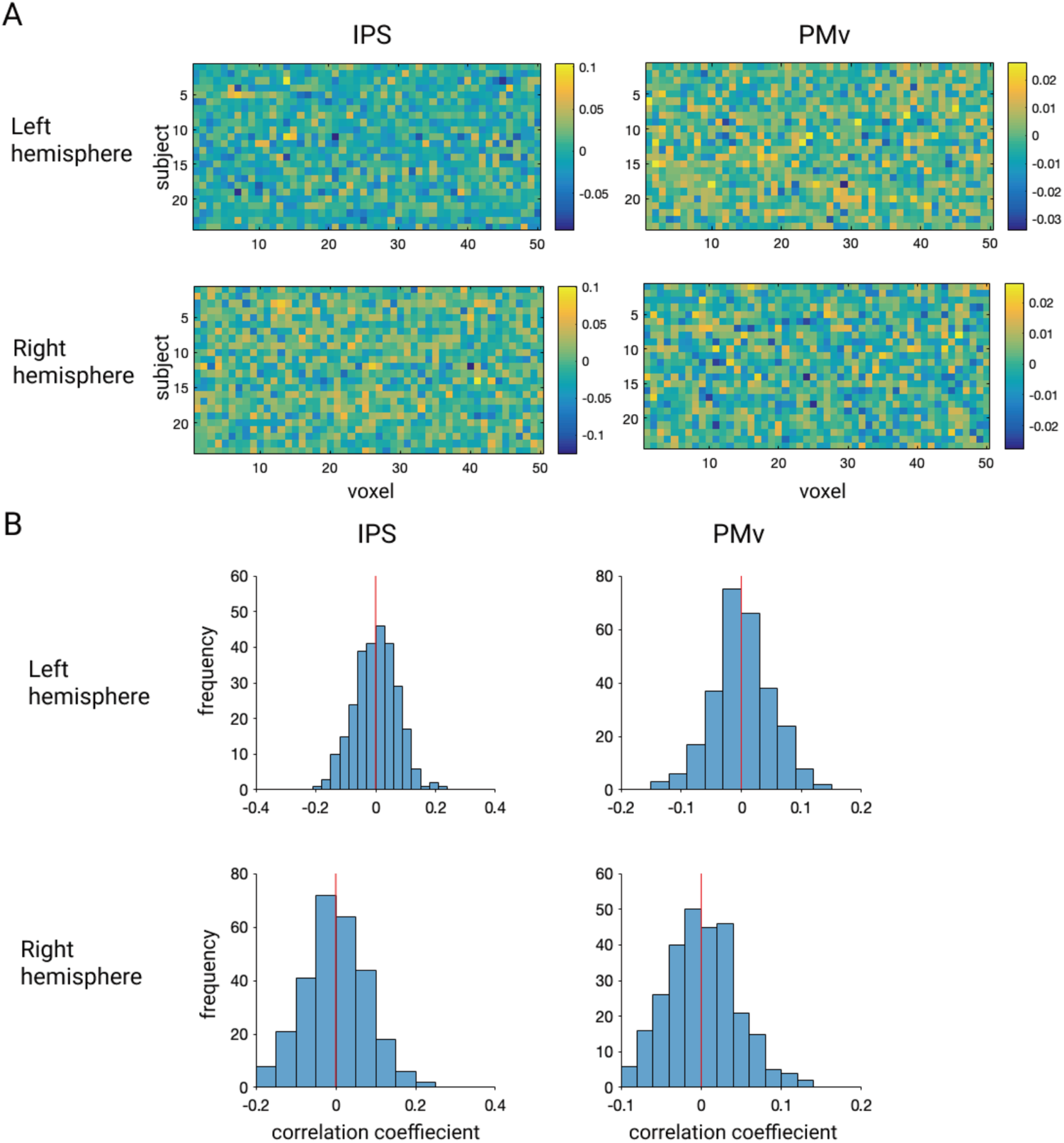
Voxel bias of classifiers. **A**, Voxel bias maps of “cross-classification” within the ROIs with significant accuracy of timings of visual feedback. **B**, Distributions of correlation coefficients (*R*) of bias between different pairs of subjects. IPS, intraparietal sulcus; PMv, ventral premotor cortex.

## Notes

### Competing Interest Statement

The authors have declared no competing interest.

